# OMICON: a community resource for studying gene coexpression networks in normal and neoplastic human brain samples

**DOI:** 10.64898/2026.08.25.747141

**Authors:** Rebecca Eliscu, Gugene Kang, Patrick G. Schupp, Daniel J. Brody, Nirmal Hariharan, Shawn Shamsian, Michael C. Oldham

## Abstract

Genome-wide coexpression analysis of intact tissue samples is a powerful approach for identifying reproducible signatures of cell types and states, since it can survey vast numbers of individuals, cells, and transcripts. However, it can be difficult to optimize gene coexpression network construction and compare results from independent analyses. To address these challenges, we developed OMICON (theomicon.ucsf.edu) for research on human brain gene coexpression networks. OMICON contains gene expression data from >17K normal and neoplastic human brain samples with standardized metadata. Systematic analysis of independent datasets identified >250K gene coexpression modules, which were characterized and compared via enrichment analysis with >40K gene sets. All modules are discoverable via an advanced search engine that can filter by genes, metadata, and enrichment results. Analyses can also be browsed with an interactive workflow visualization tool, and users can communicate within OMICON using @mention functionality to support communal research on human brain gene coexpression networks.

## INTRODUCTION

Genes do not function in isolation, but rather as coordinated programs of genomic activity that support cellular functions. Identifying these programs and understanding how they vary between normal and pathological conditions are major goals of biomedical research. We and others have shown that genome-wide coexpression analysis of intact (or ‘bulk’) human brain samples is a powerful approach for revealing highly reproducible programs (or ‘modules’) of genomic activity^1–14^, since variation in the cellular composition of bulk tissue samples inevitably drives covariation of optimal markers for cell types and cell states. Therefore, comparing bulk gene coexpression modules from different biological systems can pinpoint cell-type-specific and cell-state-specific gene expression differences without physically isolating cells^1,8,11^. This approach also provides outstanding statistical power, since it can survey gene activity representing orders of magnitude more individuals, cells, and transcripts than even the largest single-cell datasets, while eliminating many technical sources of variation that impact single-cell studies^15^.

Many methods have been developed to study gene coexpression networks. One of the best known methods is Weighted Gene Correlation Network Analysis (WGCNA), which identifies modules by clustering genes based on the dissimilarity of the topological overlap of their weighted adjacencies^16,17^. Like most clustering-based methods, WGCNA depends on user-supplied parameters that can significantly affect the number and nature of identified modules. Because it is difficult to determine which parameters are ‘best’, users must often experiment with different combinations, which can be laborious. Furthermore, it is challenging to construct a single gene coexpression network that includes all biologically ‘interesting’ modules present in a gene expression dataset, and it is difficult to discover and compare gene coexpression modules from independent studies.

To address these challenges, we have developed a flexible and iterative approach for constructing gene coexpression networks and used it to create a searchable repository of gene coexpression modules from normal and neoplastic human brain samples called OMICON (theomicon.ucsf.edu). Currently, OMICON contains >250K gene coexpression modules that were identified by analyzing 116 bulk RNA-seq and microarray datasets in the public domain, which collectively represent >17K samples from most normal adult human CNS regions and brain tumors (mostly malignant gliomas). For each dataset, we constructed up to 35 genome-wide coexpression networks by systematically altering module detection parameters, producing stereotyped outputs with novel network and module visualizations and tables listing module members and statistics. To characterize these modules, we performed enrichment analysis with >40K gene sets from the public domain. We have also created a web application called CoPA Cabana (https://oldhamlab.shinyapps.io/copacabana) that reveals the specific cellular origins of bulk coexpression modules by ‘projecting’ them onto single-nucleus RNA-seq datasets produced from normal or neoplastic human brain samples (see companion study by Kang & Oldham). Furthermore, by standardizing the metadata for all studies in OMICON, we developed an advanced search engine capable of discovering gene coexpression modules (as well as datasets, samples, and other entities) with extremely precise queries. To ensure the transparency and reproducibility of our findings, we created an interactive workflow visualization tool that provides a scaffold for inspecting and downloading results at each analysis step. Finally, to support communal interactions within OMICON, we implemented commenting functionality (including @mention functionality to direct comments to specific users) and a feedback channel for bug reports and feature requests. OMICON thus provides a novel resource to support neuroscientific and neuro-oncological research on human brain gene coexpression networks.

## RESULTS

### Data inclusion criteria

We searched for large bulk gene expression datasets produced from postnatal neurotypical human brain samples or malignant gliomas (the most common type of adult malignant brain tumor). We restricted our search to publicly available datasets produced by RNA-seq or major microarray platforms (Affymetrix, Agilent, and Illumina). We identified 49 sources of data representing 17,187 human brain samples (median = 71, range = 23 – 3,946; **Table S1**), including 10,409 normal samples, 6,615 gliomas, and a small number of other pathological samples. Select sample attributes are summarized in **Fig. 1**.

**Fig. 1.**
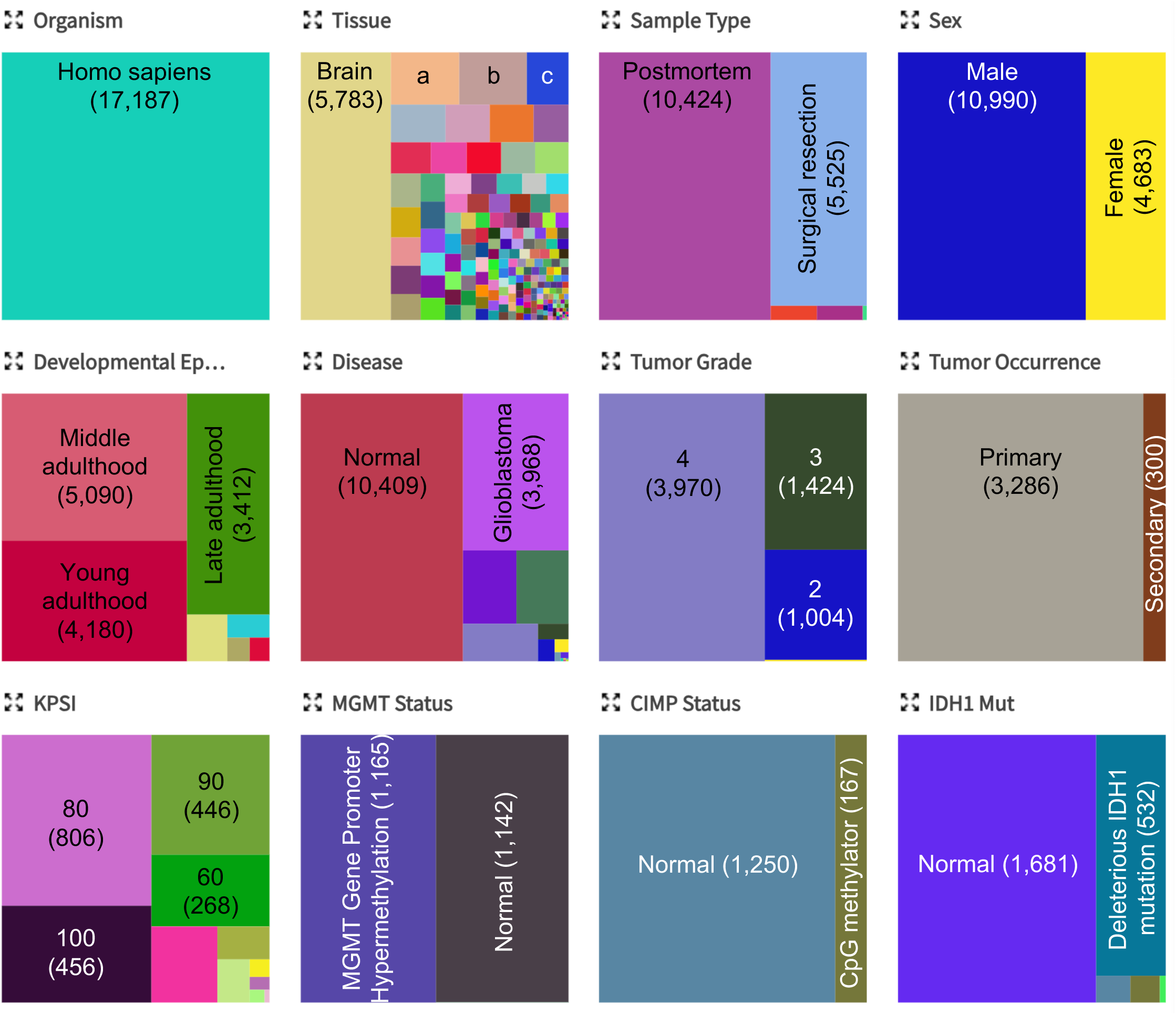
Select attributes of human brain samples in OMICON. Additional attributes are discoverable via the home page, individual *Data Collection* or *Dataset* attribute pages, or the *Advanced Search*. Parentheses: total numbers of samples for each attribute. a: frontal cortex (856). b: cerebellum (848). c: hippocampal formation (522). KPSI: Karnofsky Performance Status Index; CIMP: CpG Island Methylator Phenotype.

### Data model and metadata standardization

OMICON’s data model is summarized in **Fig. 2**. A *Data Collection* is a folder containing one or more gene expression *Datasets* from the *Data Collection* sample cohort, along with associated metadata (*Data Collection* attributes and *Sample* attributes). A *Dataset* is a folder containing a matrix of gene expression values (rows = features, columns = samples) and associated metadata (*Dataset* attributes, *Sample* attributes, and [optionally] additional *Feature* attributes). An *Analysis* is a folder containing code documents, associated metadata (*Analysis* attributes), a summary of statistics for all networks produced from the *Dataset*, and subfolders representing individual gene coexpression networks built using different parameters. Each network subfolder contains network and module visualizations and tables listing module members and statistics, along with subfolders that contain the results of gene set enrichment analyses. Additional information on these outputs is provided in the online Methods, OMICON’s online help video (‘*Gene coexpression analysis navigation*’: https://theomicon.ucsf.edu/home/help), and below.

**Fig. 2.**
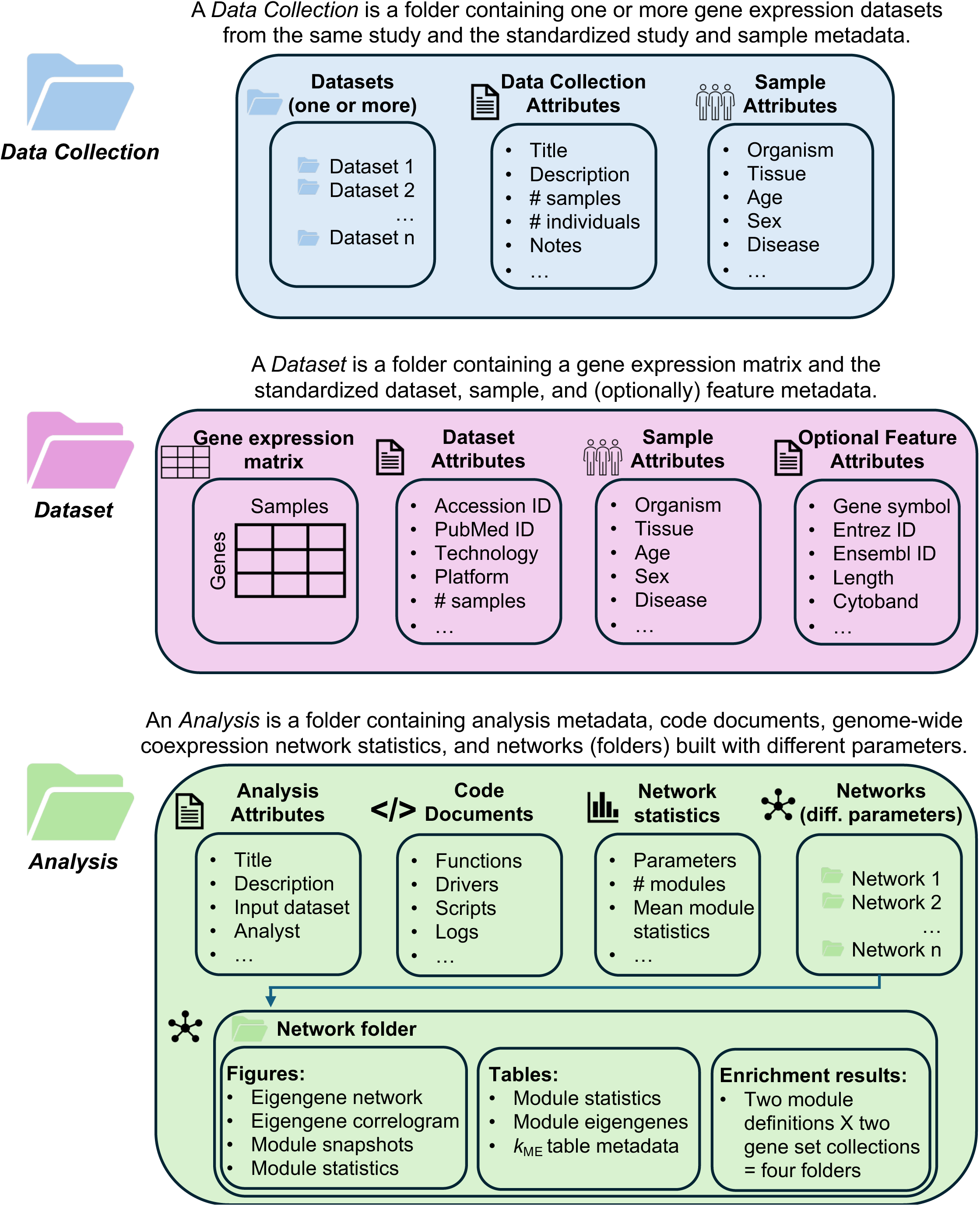
OMICON data model. A complete list of all standardized attributes is available at https://theomicon.ucsf.edu/home/help.

Comparing and integrating analysis findings from independent studies is complicated by a lack of standardized metadata. To address this challenge, we defined >125 standardized attributes for *Data Collections*, *Datasets*, *Samples*, and *Analyses* (**Table S2**). These attributes were populated using multiple biomedical ontologies, including the NCBITaxon Ontology^18^, the Uber-anatomy (UBERON) Ontology^19^, the Cellosaurus Ontology^20^, the MONDO Disease Ontology and the Human Phenotype Ontology^21^. To annotate genotypes for oncogenic mutations, we utilized identifiers from NCBI’s ClinVar database^22^, the National Cancer Institute Thesaurus^23^, and the Sequence Ontology^24^. We also defined custom attributes and values that were not adequately represented by existing ontologies and controlled vocabularies (**Table S2**). Additional information about OMICON’s data model, terminology, and metadata attributes can be found in the online help documentation (https://theomicon.ucsf.edu/home/help).

### Data analysis

Analysis of the datasets in OMICON involved three steps: i) data preprocessing, ii) genome-wide coexpression analysis, and iii) gene set enrichment analysis.

- *Data preprocessing*: Data preprocessing generally began with raw data when available. Because datasets in OMICON were produced using different technology platforms (RNA-seq and various commercial microarrays), the earliest steps of data preprocessing were variable. These steps can be reconstructed by clicking on the ‘*RDP*’ node (‘*Raw Data Processing*’) in the ‘*Workflow*’ tab (see description in *User Interface* section below) for a *Data Collection* or *Analysis* selected from the left navigation pane. Clicking on the *RDP* node reveals the code document(s) used for raw data processing, which can be previewed and downloaded. When raw data were unavailable or analysis began from processed data, the text document(s) in the *RDP* node captures information about raw data processing as performed by the original authors of the study that produced the data. Following raw data processing, data were further preprocessed using the `SampleNetwork` R function^25^, which iteratively identifies and removes outlying samples with low standardized sample connectivity, performs optional data normalization, and corrects for batch effects using the `ComBat` R function^26^. `SampleNetwork` code and outputs can be previewed and downloaded from the *OR* (‘*Outlier Removal*’), *QN* (‘*Quantile Normalization*’), and / or *BC* (‘*Batch Correction*’) nodes of a workflow.
- *Genome-wide coexpression analysis*: To enable flexible and iterative analysis of genome-wide coexpression relationships, we developed the `FindModules` R function (**Fig. 3**). This function was designed to identify the most highly correlated groups of features in a dataset. It takes as input a fully processed gene expression matrix and constructs a series of genome-wide coexpression networks based on the following user-supplied parameters (additional parameters are described in the online Methods):

- **`simType`**: This parameter defines the measure that is used to quantify genome-wide expression similarities. Supported choices include Pearson correlation, Spearman correlation, and biweight midcorrelation (‘`Bicor`’), which is a robust alternative to Pearson correlation that is less sensitive to outliers^27^. By default, all networks in OMICON use `Bicor` as the similarity measure.
- **`signumvec`**: This parameter specifies a vector of quantiles that serve as hard thresholds for gene coexpression module identification. For example, `signumvec = c(.999,.99,.98)` instructs `FindModules` to identify all groups of features whose pairwise similarities exceed the 99.9^th^, 99^th^, or 98^th^ percentiles of the genome-wide distribution of pairwise similarities for the dataset (as defined by `simType`).
- **`minsizevec`**: This parameter specifies the minimum module size. For example, `minsizevec = c(8,10,12)` instructs `FindModules` to identify all groups of at least 8, 10, or 12 features whose pairwise similarities exceed the desired similarity threshold (as defined by `simType` and `signumvec`). By default, `FindModules` will produce all possible combinations of networks defined by the `signumvec` and `minsizevec` vectors (i.e., 3 x 3 = 9 networks in the example above).
- **`merge.param`**: The dominant expression pattern of a gene coexpression module is summarized across samples by its first principal component, or *module eigengene*^16^. Because some modules may be very similar, `FindModules` uses `merge.param` to define a merging parameter. By default, modules whose eigengenes have a Pearson correlation > 0.85 are merged in an iterative fashion, starting with the pair that has the highest correlation and repeating until no pairs of modules exceed the threshold.

**Fig. 3.**
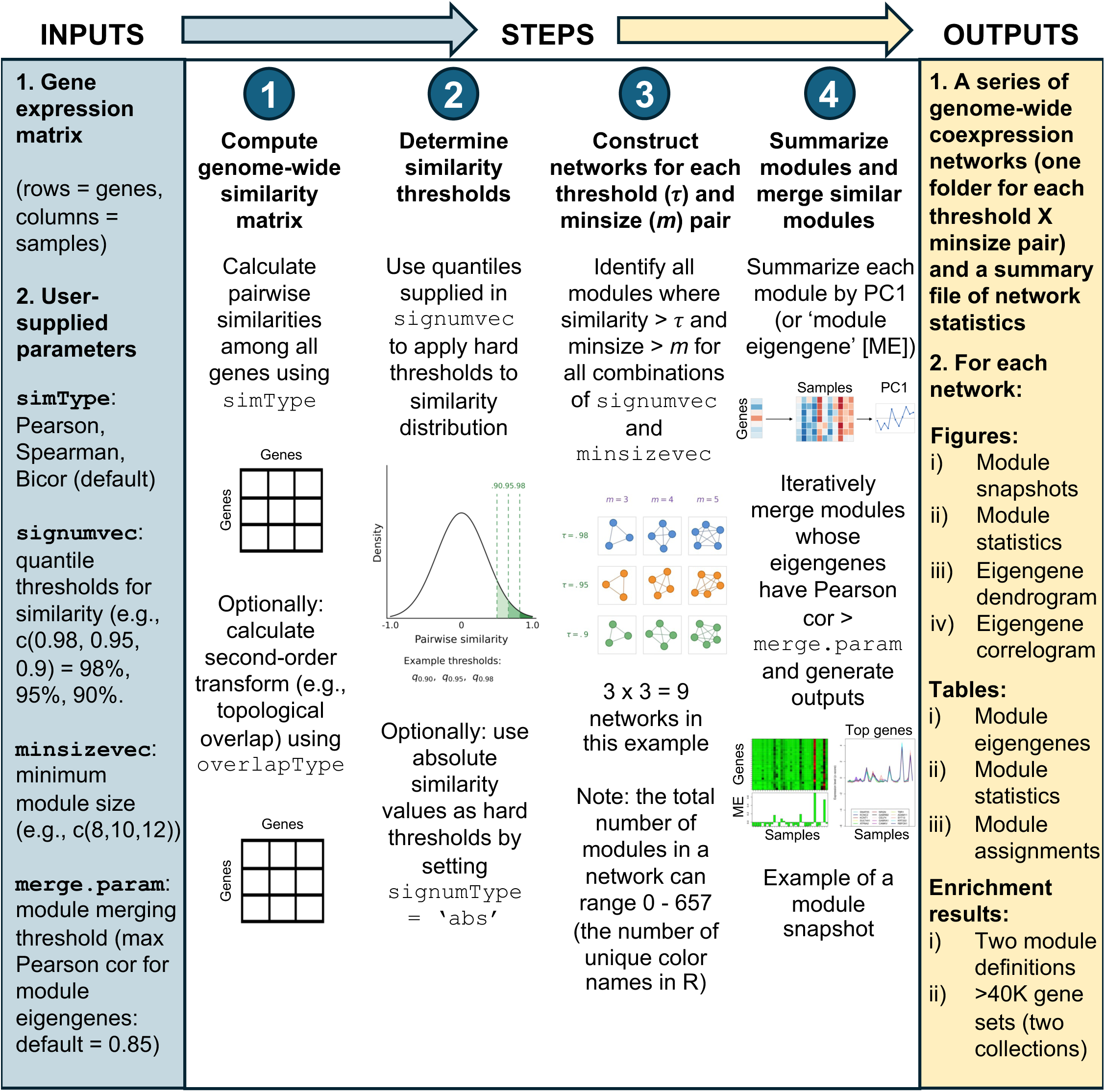
The FindModules R function.

We used `FindModules` to identify highly correlated groups of genes in normal and neoplastic human brain samples by constructing up to 35 genome-wide coexpression networks from each dataset. Each network subfolder in OMICON is named according to the parameters used to construct the network. For example, *Bicor-None_signum0.831_minSize20_merge_ME_0.85_32968* indicates that in a dataset with 32,968 features, all gene coexpression modules with group size ≥ 20 and biweight midcorrelations > 0.831 (corresponding to the 99.99^th^ percentile in this dataset) were identified and merged if the Pearson correlation of their eigengenes was > 0.85. Because each module is assigned a unique color, the number of modules is limited by the number of built-in color names in R (n = 657). If a combination of parameters yielded no modules or >657 modules, no subfolder will exist for that network.

Each network subfolder contains novel network and module visualizations and tables listing module members and statistics. For example, the network *Bicor-None_signum0.727_minSize8_merge_ME_0.85_20785* (discoverable in OMICON by searching for that folder name in the quick search bar of the masthead) identified nine highly robust gene coexpression modules in a study of malignant gliomas by Jeanmougin et al.^28^. The overall structure of the gene coexpression network is revealed by the *Module Eigengene Network* dendrogram, which is produced by hierarchically clustering the module eigengenes (**Fig. 4a**). `FindModules` also produces a *Module Eigengene Correlogram* to visualize pairwise module eigengene correlations (**Fig. 4b**).

**Fig. 4.**
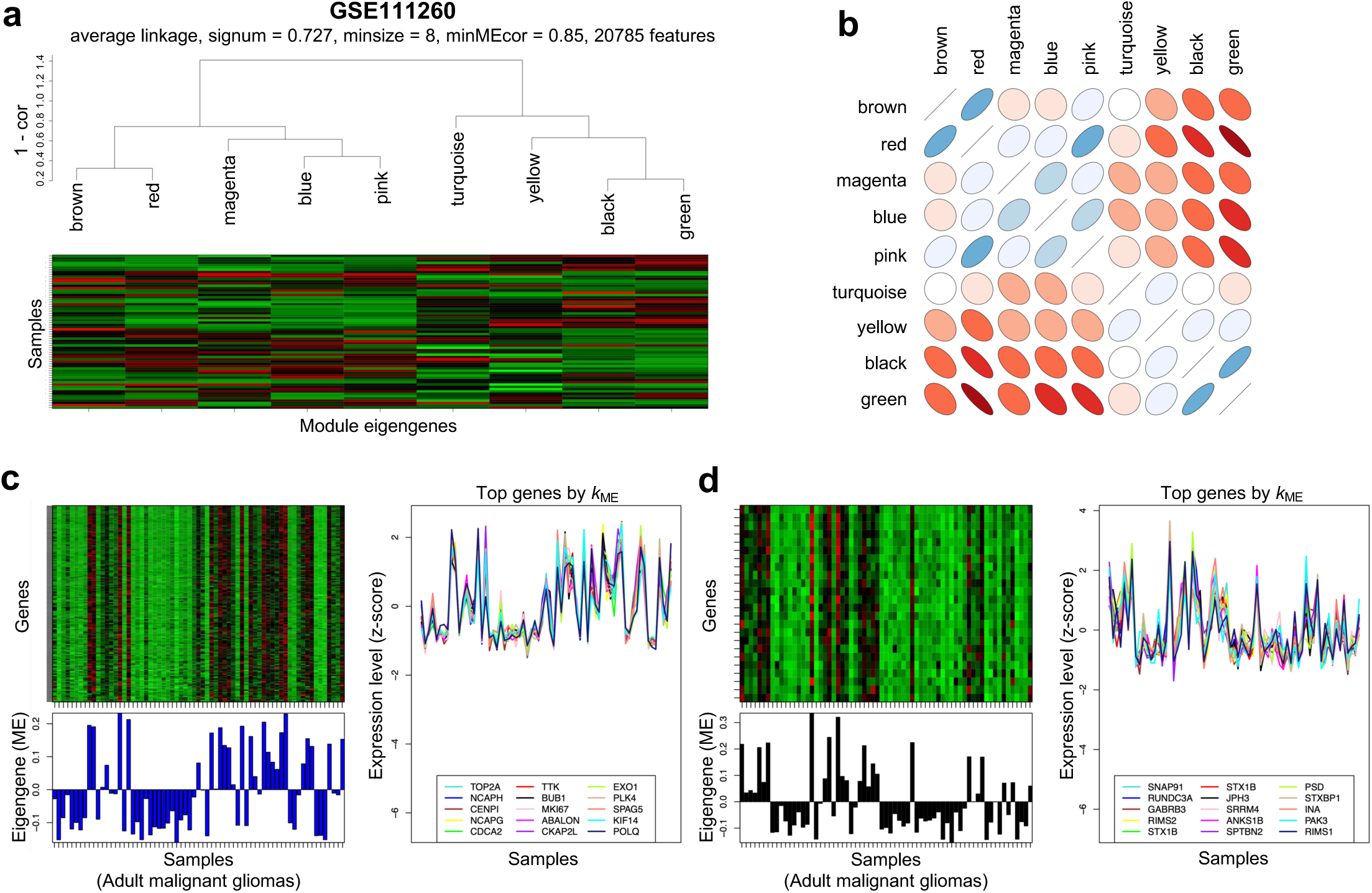
FindModules graphical outputs. Examples from analysis of GSE111260^28^. **a)** A module eigengene (ME) network produced by hierarchically clustering MEs. **b)** The corresponding ME correlogram (thin blue/red ellipse = strong +/- correlation). **c,d)** Snapshots of the blue (c) and black (d) modules, which are highly enriched with gene sets related to cell division (c) and neuronal function (d).

The gene expression patterns for the top 15 members of each module are depicted in the *Module Snapshots* file. The modules in this file are arranged in the same order as the *Module Eigengene Network* dendrogram (moving from left to right). Top members for each module are defined as those genes (or probes) with the highest Pearson correlations to the module eigengene, or *k*_ME_ values^16^. Expression values for each feature are z-scored across samples to display the cohesion of gene coexpression. For example, the blue module in this network represents a gene expression program related to cell division (**Fig. 4c**), while the black module represents a gene expression program related to neuronal function (**Fig. 4d**).

The *k*_ME_ *Table Metadata* file reports genome-wide mean expression percentiles and assignments of all genes to all modules in a network. `FindModules` provides three sets of module definitions for each network. The first (and most stringent) definition uses only the module ‘seed genes’ identified by the parameters that define the network, which are the basis for calculating the module eigengenes (‘*ModSeed*’ column in the *k*_ME_ *Table Metadata* file). The second definition includes all genes that were positively and maximally correlated with a module eigengene below a Bonferroni-corrected significance threshold (‘*TopModPosBC…*’ column in the *k*_ME_ *Table Metadata* file, where the Bonferroni correction = .05 / [# features X # modules]). The third (and most permissive) definition includes all genes that were positively and maximally correlated with a module eigengene below a false discovery rate (FDR)-corrected significance threshold^29^ (‘*TopModPosFDR…*’ column in the *k_ME_ Table Metadata* file). `FindModules` also exports a *Module Eigengenes* table that can be used for post hoc analyses (e.g., regressing module eigengenes on sample covariates), along with a table of module statistics and accompanying visualizations. For recognizability, all filenames in the same network subfolder are appended with the same suffix (e.g., ‘*10-47-01’*).

Although complete tables of *k*_ME_ values for all networks are not stored in OMICON, users can calculate these values by clicking on the vertical ellipsis (three dots) to the right of a network subfolder and selecting ‘*Create Local k_ME_ Table*’. Clicking on this link will download a zip file with instructions on how to produce the full *k*_ME_ table using an OMICON Docker image. Users can also manually create the full *k*_ME_ table by calculating the Pearson correlations between all genes and all module eigengenes using the data in the zip file.

The *Network Statistics* file in each root *Analysis* directory contains information about the parameters used to construct each network along with various network statistics. These statistics can help justify focus on particular networks depending on the user’s application. Analysis of network statistics indicates that the total number of identified modules rises quickly as the minimum module size and similarity threshold are relaxed (**Fig. S1**)^30^.

- *Gene set enrichment analysis*: To characterize gene coexpression modules, we performed enrichment analysis using two collections of gene sets from the public domain. Most gene sets are derived from the Molecular Signatures Database^31^ (“BROAD_SETS” in OMICON), which includes cytobands, pathways, Gene Ontology^32^ categories, human phenotype categories, findings from individual studies, and more. Other gene sets (“MY_SETS” in OMICON) are more neurobiologically focused and also include results from the Mouse Genome Informatics database^33^. We performed enrichment analysis for each collection of gene sets by cross-referencing them with each gene coexpression module (using *TopModPosBC…* or *TopModPosFDR…* module definitions) in each network using a one-sided Fisher’s exact test.

Enrichment analysis results for all gene sets are accessible as subfolders within each network folder and also via the *‘Enrichment results*’ nodes in *Analysis* workflows. These folders and files are named to denote the gene set collection and module definition (e.g., “GSHyperG_BROAD_SETS_10-51-14_TOPMODPOSBC”). A table includes legends and descriptions for all gene sets and enrichment P-values for all modules. For networks with ≥3 modules, an additional .PDF file extracts results for the most significantly enriched gene sets (using a Bonferroni-corrected P-value based on the # gene sets X the # modules), which are hierarchically clustered based on the dissimilarity of their enrichment P-values across all modules.

### User interface

OMICON was built in the UCSF AWS Secure Enterprise Cloud in partnership with a professional software development company (Nextware Technologies, Inc.). All data and analysis results are publicly available, but users must register to download files and post comments. From the home page, users can browse available *Data Collections* (“Data”) and *Analyses* using the left navigation pane (users can also locate specific studies by first author, accession ID, or keyword using the quick search bars). Selecting one of these opens a web page with four tabs:

- *Attributes:* This tab lists the standardized attributes of the *Data Collection* or *Analysis*. Below these attributes is a list of all *Datasets* associated with the *Data Collection* or *Analysis*, and below this list are interactive block charts of standardized sample metadata for the *Data Collection* or *Analysis* (**Fig. 1**).
- *Files*: This tab lists all files and folders that comprise the *Data Collection* or *Analysis*. Files can be previewed by clicking on the vertical ellipsis (three dots) to the right of each file. Registered users can download files and folders in the same fashion (when the download is ready, the user will be notified via a red dot over the bell icon in the masthead). Registered users can also post general questions or comments about specific files and folders by clicking on the comment icon. Comments can be directed to specific OMICON users with @mention functionality, where typing ‘@’ and the recipient’s name will direct an alert to the targeted individual about the comment. Recipients of directed comments will be notified via a red dot appearing over the bell icon in the masthead. Comments can also be added to individual *Module Snapshots* while previewing the file.
- *Workflow*: This tab provides a visual and interactive representation of the relationships between all files and folders associated with a *Data Collection* or *Analysis*. Rectangular / square nodes denote data containers (nouns), while circular nodes denote code transformations (verbs), providing a precise narrative structure for reproducible analysis of each dataset. Hovering over a node will reveal a pop-up listing its attributes and files, which can be previewed and downloaded (**Fig. 5**).
- *Comments*: This tab allows users to post comments or questions about a *Data Collection* or *Analysis*. All comments in OMICON (including comments that have been directed with @mention functionality) are visible to all users.

**Fig. 5.**
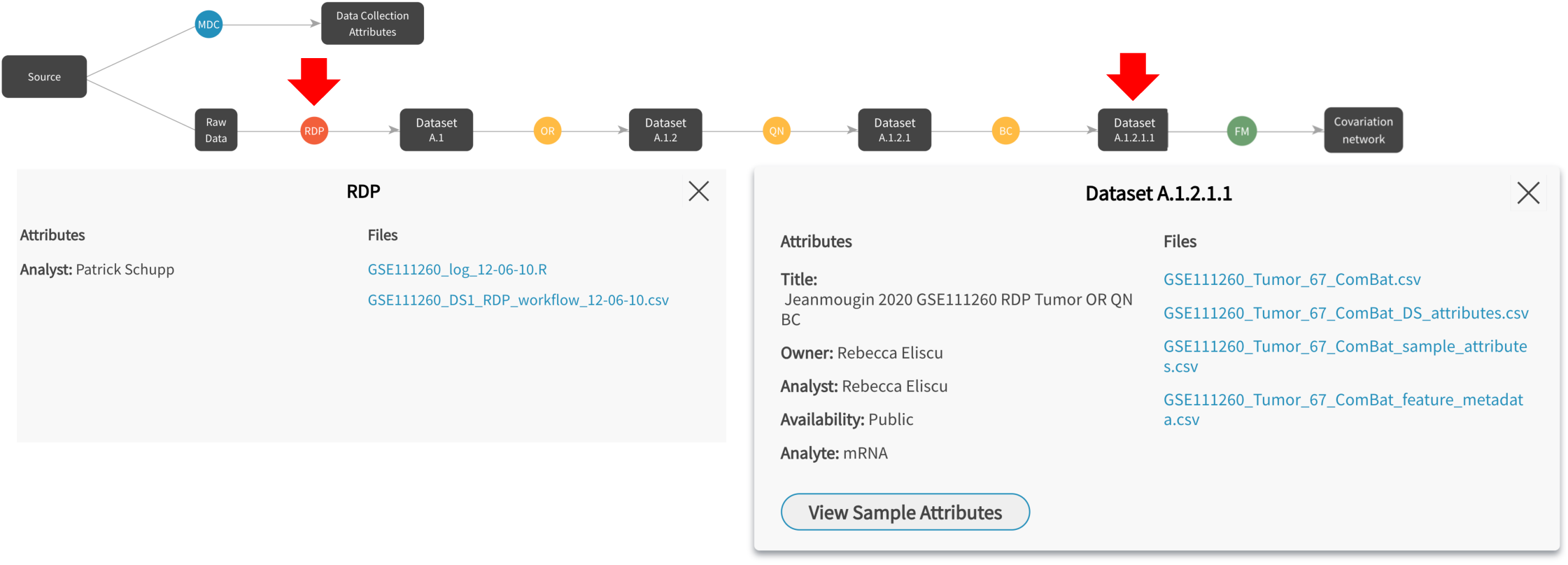
Interactive workflow visualization tool. Clicking on the ‘Workflow’ tab of any Data Collection or Analysis will reveal the precise steps involved in the workflow. Black rectangles are standardized data packages and colored circles are code transformations. Hovering over a node will reveal its attributes and associated files for preview and download (examples: red arrows). This tool ensures provenance and reproducibility for all results in OMICON. MDC = make data collection; RDP = raw data processing; OR = outlier removal; QN = quantile normalization; BC = batch correction; FM = `FindModules`.

### Additional resources

- Clicking on *‘Resources*’ in the left navigation pane will direct registered users to more information about OMICON’s sample and feature annotations:

- *Sample annotations*: This section provides links to the primary ontologies and controlled vocabularies used to annotate biological samples in OMICON.
- *Feature annotations*: This section provides links to resources that uniquely identify genes, gene products, or gene sets. Any of these identifiers can be used to search for gene coexpression modules (by selecting the *‘Feature Input Type*’ under ‘*Advanced Search*’ → ‘*Filter Data*’ → ‘*Feature Attributes*’). Genome-wide relationships between these identifiers are also provided in the ‘*Mapping Tables*’ sub-section, which includes mapping tables for microarray datasets (‘*Microarray Mapping Tables*’) and RNA-seq datasets (‘*Organism Mapping Tables*’). This section also includes a searchable repository of all gene sets known to OMICON (n=41,421) and their associated metadata (gene set name / IDs, source, category, size, species, PubMed ID, and description). By clicking on ‘*Gene Sets*’, users can search for gene sets of interest using keywords in the quick search bar (e.g., ‘astrocytes’). By clicking on the ‘*Advanced Search*’ link in the *Gene Sets* resource, users can filter gene sets by species, category, source, gene symbols of interest (comma-separated), and gene set size. Individual gene sets can be previewed and downloaded by clicking on the ellipsis next to each *Set ID*.

- *System Tables:* Clicking on the gear icon located on the right side of the masthead will produce links to system tables that define the metadata (attributes) for *Data Collections*, *Datasets*, *Samples*, and *Analyses*.
- *Notifications:* Clicking on the bell icon in the masthead will let users view their notification history for searches, downloads, and comments.
- *Help documentation:* Clicking on ‘*Help*’ in the masthead provides links to recorded videos describing OMICON’s functionality (‘*Overview*’, ‘*Data navigation*’, ‘*Gene coexpression analysis navigation*’, ‘*Other resources*’, and ‘*Advanced search*’). There is also a glossary of terminology that defines all objects and attributes under OMICON’s data model.
- *Feedback:* Clicking on ‘*Feedback*’ in the masthead produces a form for users to submit bug reports and feature requests. To submit a bug report or feature request, users must supply their name, email, and a brief description of the issue. Completing this form automatically generates a ticket in Jira project management software for review and prioritization by the development team.

### Advanced search

By standardizing the metadata for all data and analysis results in OMICON, we developed an advanced search engine capable of extremely precise queries. The *Advanced Search* (accessible from the masthead link) provides a flexible interface to identify data, analysis products, or file types with desired attributes. Users must first select one of these primary result types (**Fig. S2a**). If users select ‘*Data*’ or ‘*Analyses*’, they must also specify which data or analysis product they are searching for along with additional attributes to return with the search results using the check boxes to the right (**Fig. S2b,c**). Under ‘*Data*’, users can search for *Data Collections*, *Datasets*, *Samples*, or *Features* (i.e., genomic features). Under ‘*Analyses*’, users can search for *Analysis Instances*, *Covariation Networks* (groups of *Covariation Modules* identified in the same dataset with the same parameters), *Covariation Modules* (groups of genomic features with similar patterns in a dataset), and *Gene Sets*. For each result subtype, users can apply conditions (filters) located on the ‘*Data*’ and ‘*Analyses*’ sub-tabs to restrict the search results (**Fig. S2d,e**). The use of AND / OR logic and grouping of conditions enables highly specific search queries, as illustrated by the examples in the ‘*Sample Queries*’ section of the Advanced Search. By default, OMICON will retain a user’s 30 most recent queries (*Advanced Search* → *Recent Queries*). Users can also save the structure and results of a query by clicking on the ‘*Save query & search results*’ button that is provided with the search results (**Fig. S2f**). Each saved query receives a unique URL that can be shared with other users. Finally, users can also add comments to saved queries using the ‘*Add comments*’ button that is provided with the search results. Additional details on search functionality are provided in the online help video (‘*Advanced search*’).

### Use cases

By standardizing and organizing gene coexpression analysis of normal and neoplastic human brain transcriptomes representing vast numbers of individuals, cells, and transcripts, OMICON supports several use cases:

- *Browsing gene coexpression networks for datasets of interest*: OMICON makes it easy to browse the most highly correlated groups of genes in a given dataset. By clicking on ‘*Analyses*’ in the left navigation pane, users can select a dataset of interest and browse available gene coexpression networks by clicking on the ‘*Files*’ tab, which lists networks (folders) from the smallest to the largest number of modules, or by clicking on a ‘*Covariation network*’ container on the ‘*Workflow*’ tab, which lists network attributes and network files. Within a network folder or container, users can examine gene coexpression modules by previewing the *Module Snapshots* file (named “Modules_<<TIMESTAMP>>.pdf”). Alternatively, users can search for gene coexpression networks with desired attributes via the *Advanced Search*.
- *Discovering gene coexpression modules with desired attributes*: The *Advanced Search* enables highly specific and flexible queries to identify gene coexpression modules with desired attributes. Users can search for modules that are significantly enriched (below a specified P-value threshold) with a gene set of interest (selecting from >40K gene sets via the ‘*Gene Sets*’ search interface: https://theomicon.ucsf.edu/home/resources/gene-set). Users can also perform queries to identify modules that are significantly enriched with all gene sets whose names include a particular keyword (e.g., ‘microglia’, as illustrated in sample query S7 in the *Advanced Search*). In addition, users can require that modules contain one or more comma-separated genes of interest (where genes can be specified by gene symbols or other unique identifiers, including Entrez ID, RefSeq ID, Ensembl ID, UniProt ID, and more – see list of ‘*Feature Input Types*’ on the ‘*Feature Attributes*’ tab of *Advanced Search*). For example, **Fig. 6** shows snapshots of modules identified with the following parameters (https://theomicon.ucsf.edu/home/advanced-search?queryId=c9c4d54cc0ad4106bc9a761e450dbf35): i) significant enrichment (P < 1e-10 using the *TopModPosBC* [Bonferroni] module definition described above) with gene set MOSET7037, which represents a list of published T cell markers^34^); ii) presence of >2 of the following known T cell markers as module seed genes: *CCL5*, *CD2*, *CD3D*, *CD3E*, *CD3G*, *CD8A*, *GZMA*, *GZMK*, *LCK*; and iii) derived from datasets where 100% of samples are from glioblastomas (**Table S1**). This query therefore identifies highly reproducible gene coexpression signatures of human T cells in the GBM microenvironment driven by vast numbers of cells.
- *Enabling downstream analyses:* The *Advanced Search* can also be used to identify modules or datasets for downstream analyses. One such analysis involves quantifying relationships between modules of interest and sample covariates, which is often performed via linear modeling of module eigengenes. Another type of analysis involves aggregating genome-wide *k*_ME_ vectors for similar modules from independent datasets to identify optimal markers of cell types or states at scale^11^. A third type of analysis seeks to identify gene coexpression relationships that are present in one dataset but not another through differential coexpression analysis (**Fig. 7**).
- *Integration with single-cell data*: Integrating bulk gene coexpression modules with single-cell datasets can clarify the cellular origins of highly reproducible patterns of gene activity. In a related study by Kang & Oldham (2026), we describe a novel approach to perform this integration called **Co**variation **P**rojection **A**nalysis (CoPA). To apply CoPA to gene coexpression networks in OMICON, we have created a web application called CoPA Cabana (https://oldhamlab.shinyapps.io/copacabana) where users can upload *k*_ME_ Table Metadata (i.e., module assignments for a given gene coexpression network) from OMICON (**Fig. 8a,b**). Absolute and relative expression levels of genes comprising each module are then summarized in the selected single-cell atlas(es) for all annotated cell types. For example, the turquoise module featured in **Fig. 8c**, which is highly enriched with gene sets related to cell division, is primarily expressed by dividing neoplastic cells in the CELLxGENE^35^ Core GBmap (https://cellxgene.cziscience.com/e/c888b684-6c51-431f-972a-6c963044cef0.cxg/), as expected (**Fig. 8d**). However, at the most granular annotation level we also see evidence of expression by other proliferative populations, including T cells and smooth muscle cells. This strategy provides a novel and powerful approach for integrating vast quantities of bulk and single-cell data.

**Fig. 6.**
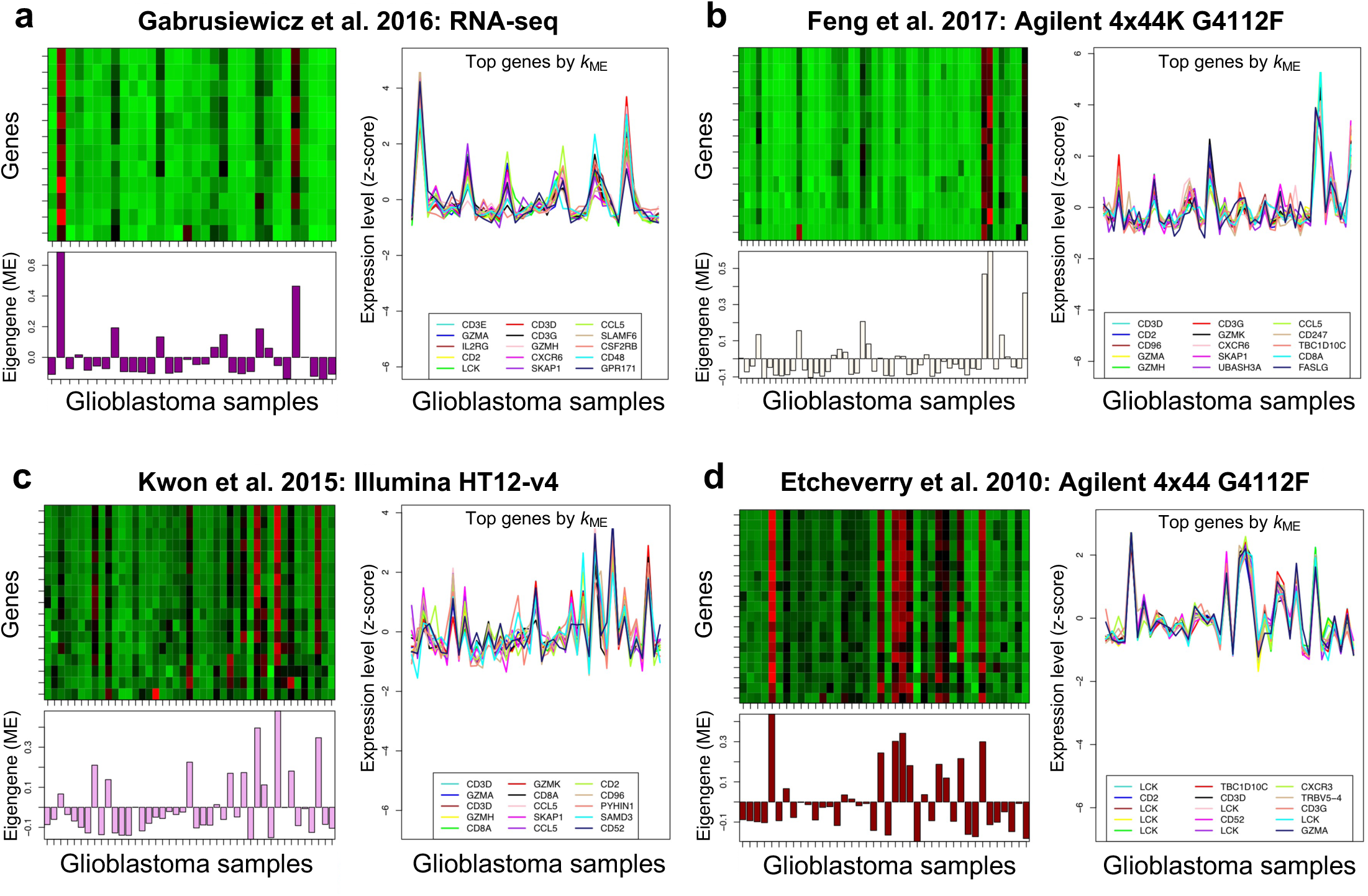
OMICON enables discovery of gene coexpression modules with desired attributes. **a-d)** Representative gene coexpression signatures of human T cells in the glioblastoma microenvironment identified by OMICON’s *Advanced Search* (see query description and link in main text).

**Fig. 7.**
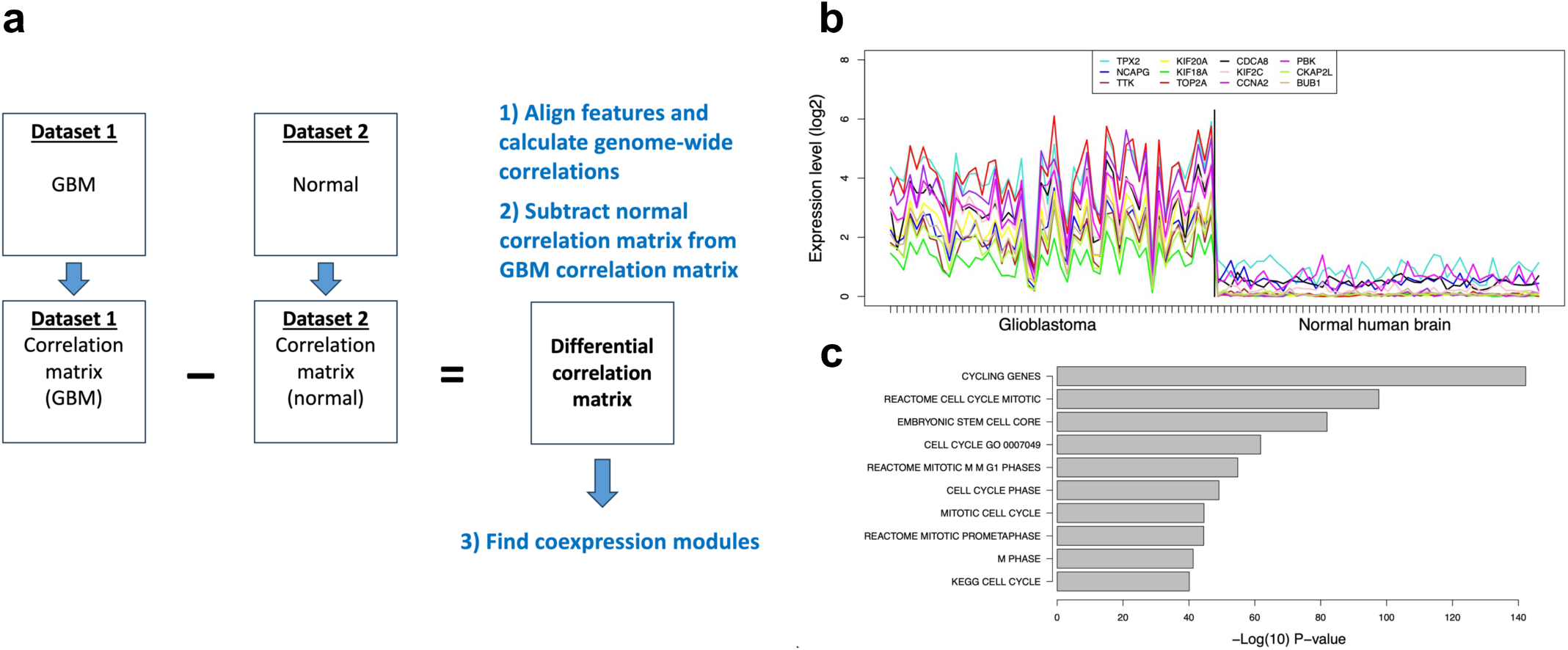
OMICON facilitates diverse downstream applications such as differential coexpression analysis. **a)** Strategy for identifying gene coexpression modules that are present in glioblastoma (GBM) but absent in normal human brain. **b-c)** Example of one such module, which is significantly enriched with gene sets related to cell division. GBM dataset: Clinical Proteomic Tumor Analysis Consortium (<u>OMICON.DS.001395.A.1.1.2.1.1</u>); normal human brain dataset: Allen Brain Institute (<u>OMICON.DS.001445.A.1.1</u>) (**Table S1**).

**Fig. 8.**
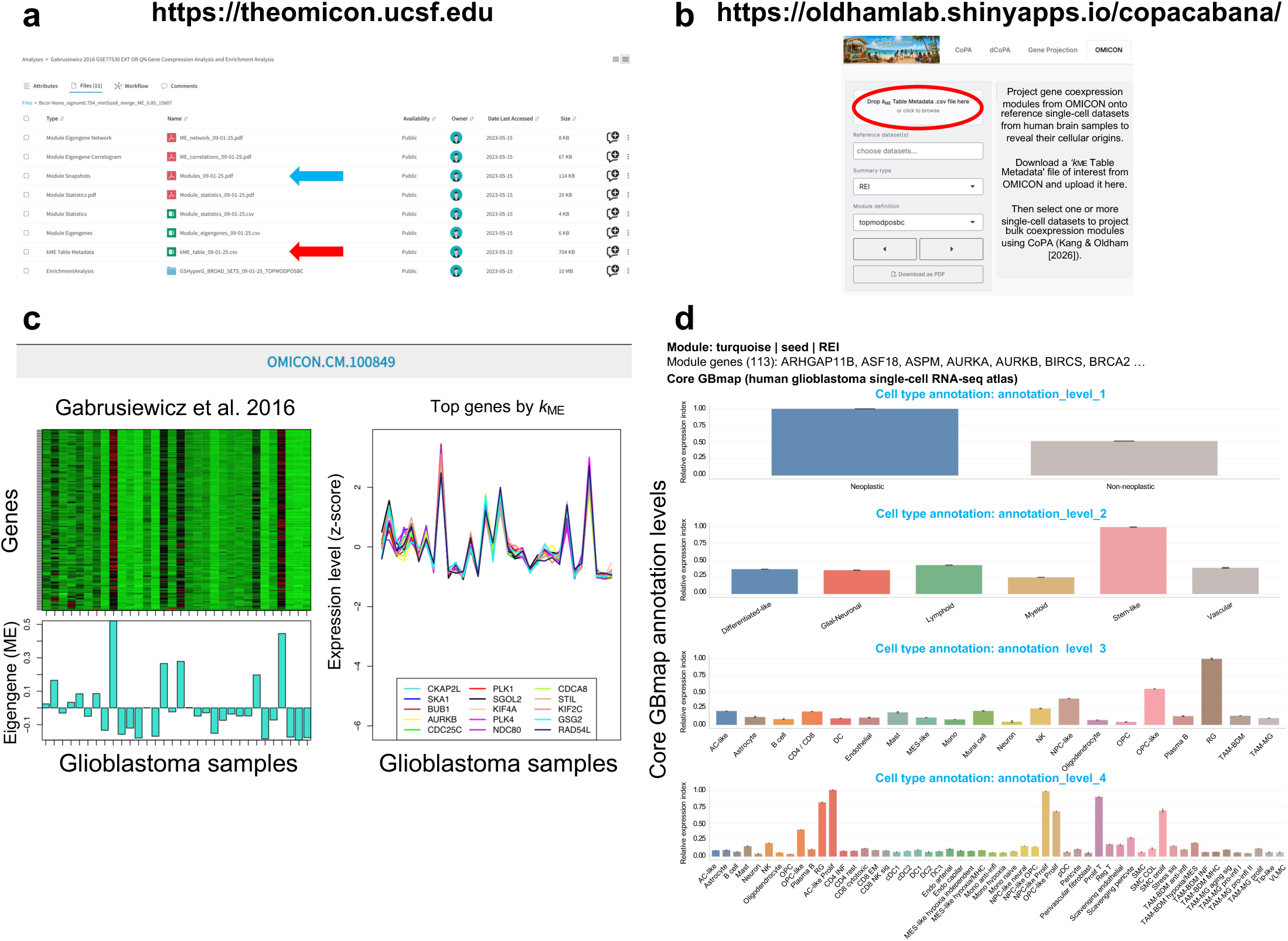
Covariation Projection Analysis (CoPA) on the CoPA Cabana web site integrates bulk gene coexpression modules with single-cell analysis. **a)** Step 1: Download module assignments for a covariation network of interest from OMICON (*k*_ME_ Table Metadata file: red arrow). **b)** Step 2: Upload the *k*_ME_ Table Metadata .csv file to CoPA Cabana (red circle). **c)** Identify one or more modules of interest in the Module Snapshots file (turquoise arrow in **a**). **d)** Select the single-cell dataset(s) and module of interest from the pulldown menus. Expression levels of all module genes are averaged and normalized to mean genome-wide expression levels to calculate a relative expression index for each cellular population. See companion study by Kang & Oldham (2026) for additional details.

## DISCUSSION

We have developed OMICON to foster communal research on gene coexpression networks in normal and neoplastic human brain samples. By identifying groups of genes with the strongest correlations in large bulk datasets, we have revealed highly reproducible modules of genomic activity derived from thousands of individuals, billions of cells, and hundreds of billions of transcripts. Through standardization of metadata and analysis pipelines, including module enrichment analysis with >40K gene sets, we have provided a rich and searchable set of results to focus neuroscientific discourse and generate novel hypotheses about gene activity in normal and neoplastic human brain samples. Furthermore, we have produced a companion web site called CoPA Cabana (https://oldhamlab.shinyapps.io/copacabana) that reveals the cellular origins of bulk coexpression modules by ‘projecting’ them onto single-cell RNA-seq datasets produced from normal or neoplastic human brain samples (see companion study by Kang & Oldham). The development of OMICON highlights several important scientific and meta-scientific issues.

Genome-wide coexpression analysis of bulk tissue samples is a powerful approach for surveying the population structure of gene activity in biological systems. Inevitably, variation in the abundance and activity of cell types and states drives coexpression of genes most strongly associated with those cell types and states^11^. We have shown how aggregating similar gene coexpression modules in bulk datasets produced from the same biological system can reveal optimal markers of cell types and states at scale, while comparing similar modules in different biological systems can pinpoint cell-type-or cell-state-specific gene expression differences without physically isolating cells^1,8,11,13^. This strategy provides enormous statistical power for target discovery and complements the use of single-cell assays for studying individual genes.

OMICON addresses two important meta-scientific challenges that impede biomedical research. The first issue is the lack of metadata standardization across studies, which creates inefficiencies by hampering the discovery, interpretability, interoperability, and reproducibility of published data and analyses^36–38^. To address this challenge, we leveraged biomedical ontologies and controlled vocabularies to standardize metadata, which enabled development of an advanced search engine for discovering data and analysis results with desired attributes. The second challenge is the difficulty reproducing large-scale meta-analyses described in published studies, which may involve huge amounts of data, code, and intermediate analysis products. To address this challenge, we developed an interactive workflow visualization tool to provide a scaffold for inspecting and downloading data and code at each step of our analyses, thereby ensuring transparency and reproducibility. These solutions provide proofs of concept that can be generalized for other types of data and applications in the future.

OMICON has several limitations. Although the `FindModules` R function thoroughly samples the high-correlation space in each dataset, the default module merging parameter may combine similar patterns of gene activity derived from different cellular sources. For example, the transcriptomes of most neuronal subclasses are very similar (genome-wide r ≍ 0.9; see Kang & Oldham), which makes it difficult to isolate coexpression modules driven by a specific neuronal subclass; on the other hand, integrative analysis of bulk and single-nucleus RNA-seq data suggests that such modules may be very rare (see Kang & Oldham). Although OMICON does not yet store single-cell or single-nucleus datasets or analysis results, we have developed a web application where users can project bulk coexpression modules from OMICON onto single-cell or single-nucleus RNA-seq datasets from normal and pathological human brain samples to reveal their cellular origins (https://oldhamlab.shinyapps.io/copacabana). In the future, we will extend these efforts by assimilating gene expression datasets and coexpression networks from other normal and pathological human tissue samples, developing an API to support programmatic access, and extending our methodology to other types of omics datasets.

In summary, by harnessing the statistical power of bulk tissue sampling, OMICON provides a novel resource for studying gene coexpression networks in normal and neoplastic human brain samples. We welcome suggestions for improvement via OMICON’s *Feedback* channel and hope that this resource will promote dialogue and discoveries that lead to novel treatments for neuropathologies.

## METHODS

### OMICON solution architecture

A high-level overview of OMICON’s solution architecture is shown in **Fig. S3**. The OMICON platform supports three distinct user roles: guest users for basic data exploration, registered users with full capabilities, and internal members responsible for data management and administrative functions. The system adopts a modern layered architecture where a React-based frontend communicates with a Django backend via secure APIs. The backend is responsible for managing logic, controlling user access, and coordinating search operations. The entire platform is hosted on AWS, ensuring enhanced security, scalability, and reliability. Data are managed through a hybrid storage approach: a relational database handles users, metadata, ontology, and collaboration data, while a search engine enables fast and efficient querying of datasets, samples, and analysis results. The application supports real-time and asynchronous processing. Simple searches provide instant results, while complex queries are handled in the background, enabling users to continue work without interruption. Users are notified once results are available. Additionally, the platform allows registered users to preview and download data files, save search results for future use, and collaborate through comments, providing a collaborative solution for transcriptomic data exploration and analysis.

### Data analysis

All data analysis steps can be reconstructed using OMICON’s interactive workflow visualization tool, which is available on the ‘Workflow’ tab for each *Data Collection* and *Analysis* in OMICON. In this representation, rectangular / square nodes denote data containers and circular nodes denote analysis steps (hovering over a node reveals a pop-up listing its attributes and files, which can be previewed and downloaded). The major analysis steps are as follows:

- *Raw Data Processing (RDP):* Clicking on the RDP node reveals the code document(s) used for raw data processing. This node may also contain a table that summarizes the different steps of the RDP workflow. If raw data were unavailable or analysis began from processed data, the *RDP* node captures information about how the authors of the study that produced the data performed raw data processing.
- *Outlier Removal (OR), Quantile Normalization (QN), and Batch Correction (BC):* Clicking on any of these nodes reveals the code documents used by the `SampleNetwork` R function, which performs all of these data preprocessing steps in succession^25^. These nodes also contain intermediate files and figures produced by `SampleNetwork`, which help inform user decisions on appropriate data preprocessing choices. `SampleNetwork` also calls the `ComBat` R function to correct for batch effects as needed^26^.
- *FindModules (FM)*: Clicking on an FM node reveals the code documents and parameters used by the `FindModules` R function to identify gene coexpression modules. Each FM node therefore produces a unique genome-wide coexpression network built using different parameters. The outputs of a `FindModules` run are stored in the *Covariation network* node immediately downstream of the FM node.
- *Enrichment Analysis (EA)*: Clicking on an EA node reveals the driver code used to perform module enrichment analysis with two large collections of gene sets derived from the Molecular Signatures Database^31^ (“BROAD_SETS”) and the Oldham lab (“MY_SETS”). For each gene set collection, EA was performed twice: i) using all genes that were positively and maximally correlated with a module eigengene (PC1 of a coexpression module) below a Bonferroni-corrected significance threshold (“TOPMODPOSBC”), or ii) an FDR-corrected significance threshold (“TOPMODPOSFDR”). The outputs of each EA are stored in the *Enrichment results* node immediately downstream of the EA node.
- *Make Data Collection (MDC)*: Clicking on the MDC node reveals housekeeping code and accessory files used to standardize metadata and package files for a *Data Collection*. The output of MDC includes standardized *Data Collection* and *Sample* attributes, which are stored in the *Data Collection Attributes* node immediately downstream of the MDC node.

### Web site information

OMICON is accessible at https://theomicon.ucsf.edu. It is free to use but requires registration to download files and post comments. Users can also analyze the cellular origins of bulk gene coexpression modules in OMICON by downloading the *k*_ME_ *Table Metadata* file from a *Covariation Network* folder, uploading it to a custom R Shiny app at https://oldhamlab.shinyapps.io/copacabana/, and selecting one or more single-cell or single-nucleus RNA-seq datasets for module projection analysis as described in the companion study by Kang & Oldham (2026).

## ACKNOWLEDGMENTS

We are grateful to Mattice Harris (UCSF) and other members of UCSF IT for technical support. We thank Marlene Grenon (UCSF) and Randi Jenkins (UCSF) for helpful discussions, as well as other Oldham lab members for feedback and usability testing. We also thank Nextware Technologies team members Sanoj Sudhakaran, Shravan Suresh, Pratheeksh Joseph, Gouri Lekshmi, Shiny Raj, Aswathy Vijay, Sharat Nair, Alex Jacob, Vineet Ravindran, Mizba Feroz, Akshay Ajayakumar, Renjith Reghuvaran, Dilip Kumar, Subin Babu, Vismay Valsara, Muneeb Ali, Remya Vava, Carolina Escobar, Divya Padmanabhan, and Martin Thormann for their efforts to develop the OMICON application. This work was supported by NIH/NIMH R01 MH113896, NIH/NCI R01 CA244621, NIH/NIMH R01 MH123156, NIH/NCI R01 CA292649, NIH/NCI SPORE Developmental Research Projects, and the Carl Kawaja Gift Fund (M.C.O.).

## AUTHOR CONTRIBUTIONS

M.C.O. conceived OMICON and wrote most of the code for data analysis. R.E. optimized code, performed most data analyses, and prepared data for ingestion. M.C.O. and R.E. standardized metadata and prepared the manuscript contents for feedback from other authors. G.K. created the CoPA Cabana R Shiny app to project OMICON gene coexpression modules onto single-cell RNA-seq datasets. R.E., G.K., P.G.S., and D.J.B. performed usability testing, conducted additional analyses, and provided suggestions for application improvement. M.C.O., N.H., and S.S. designed application functionality and S.S. served as overall software development lead for the OMICON application.

## DATA AND CODE AVAILABILITY

All gene expression datasets analyzed in this study are publicly available (**Table S1**). As described in the manuscript, users can download datasets directly from OMICON using the *Files* or *Workflow* tabs, or via links to their data repository accession IDs (which are also stored in OMICON). Code used to analyze gene expression datasets can also be downloaded directly from OMICON by navigating to the *Workflow* tab for a given *Data Collection* or *Analysis* and clicking on the circular icon representing a particular analysis step, as described in the manuscript.

## EXTENDED DATA FIGURE LEGENDS

**Fig. S1.**
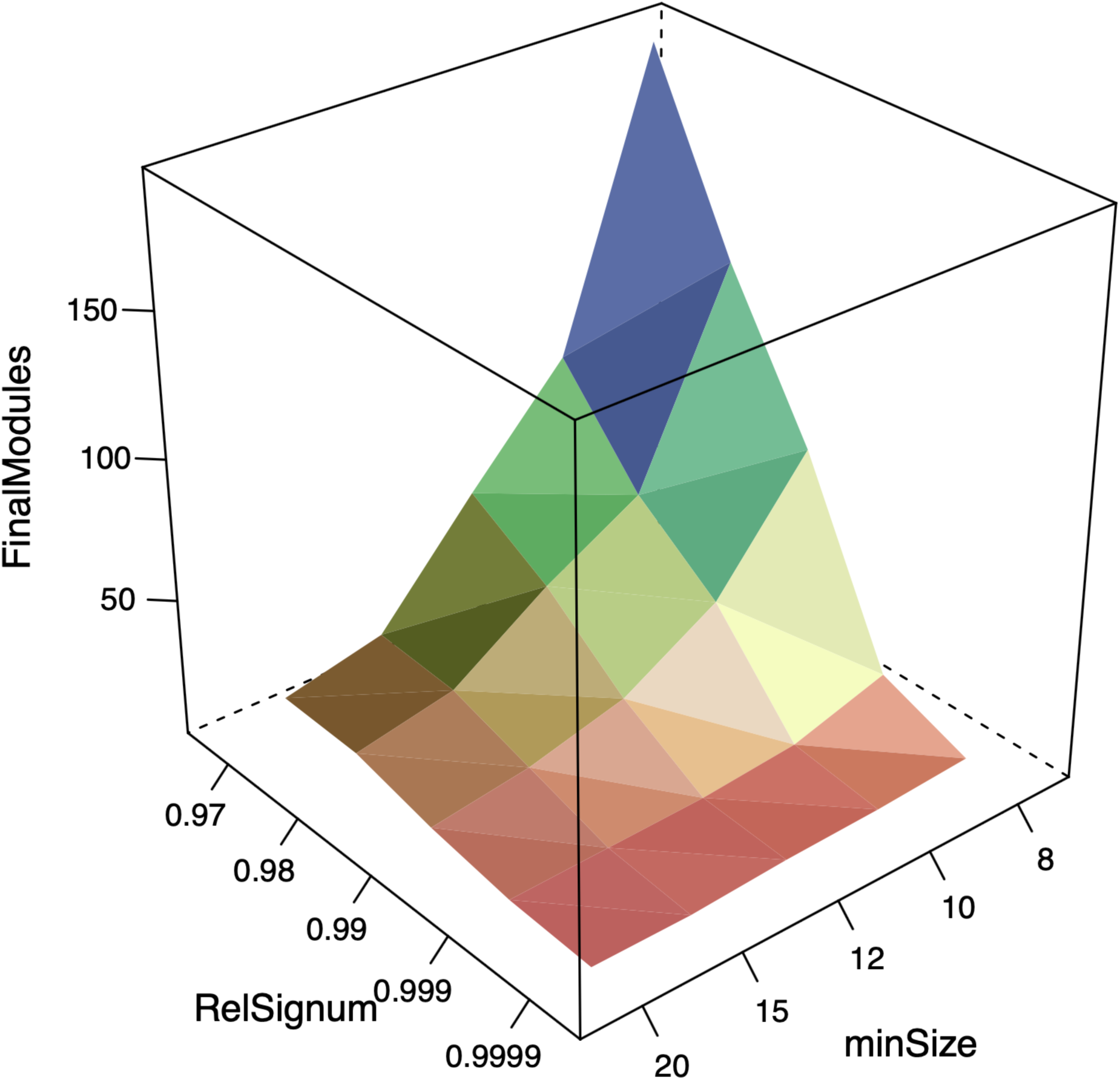
The number of modules in a network rises quickly as the minimum module size and similarity threshold are relaxed. Representative relationships between hard thresholds (quantiles), minimum module size, and the total number of identified modules. Analysis performed on malignant glioma samples from GSE4290^30^.

**Fig. S2.**
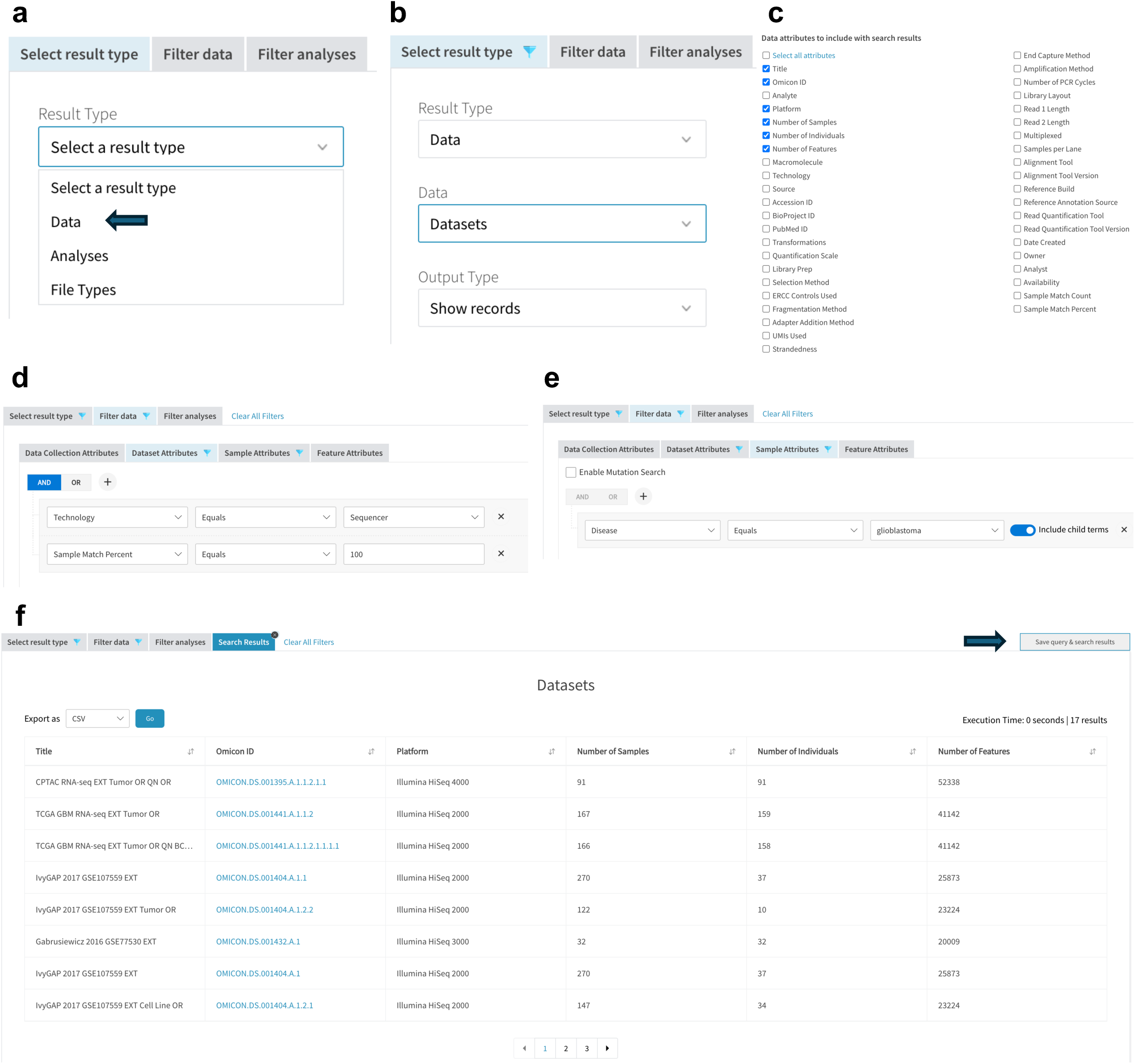
Advanced Search. **a)** Users can search for data products, analysis products, of file types in OMICON. **b)** Here, the user selects Datasets as the result type. **c)** Tabular outputs can be configured by selecting column headers to include. **d-e)** Dataset and Sample attributes are selected on separate subtabs. **f)** A table of linked datasets matching search criteria, which can be saved (arrow) and shared with a unique URL.

**Fig. S3.**
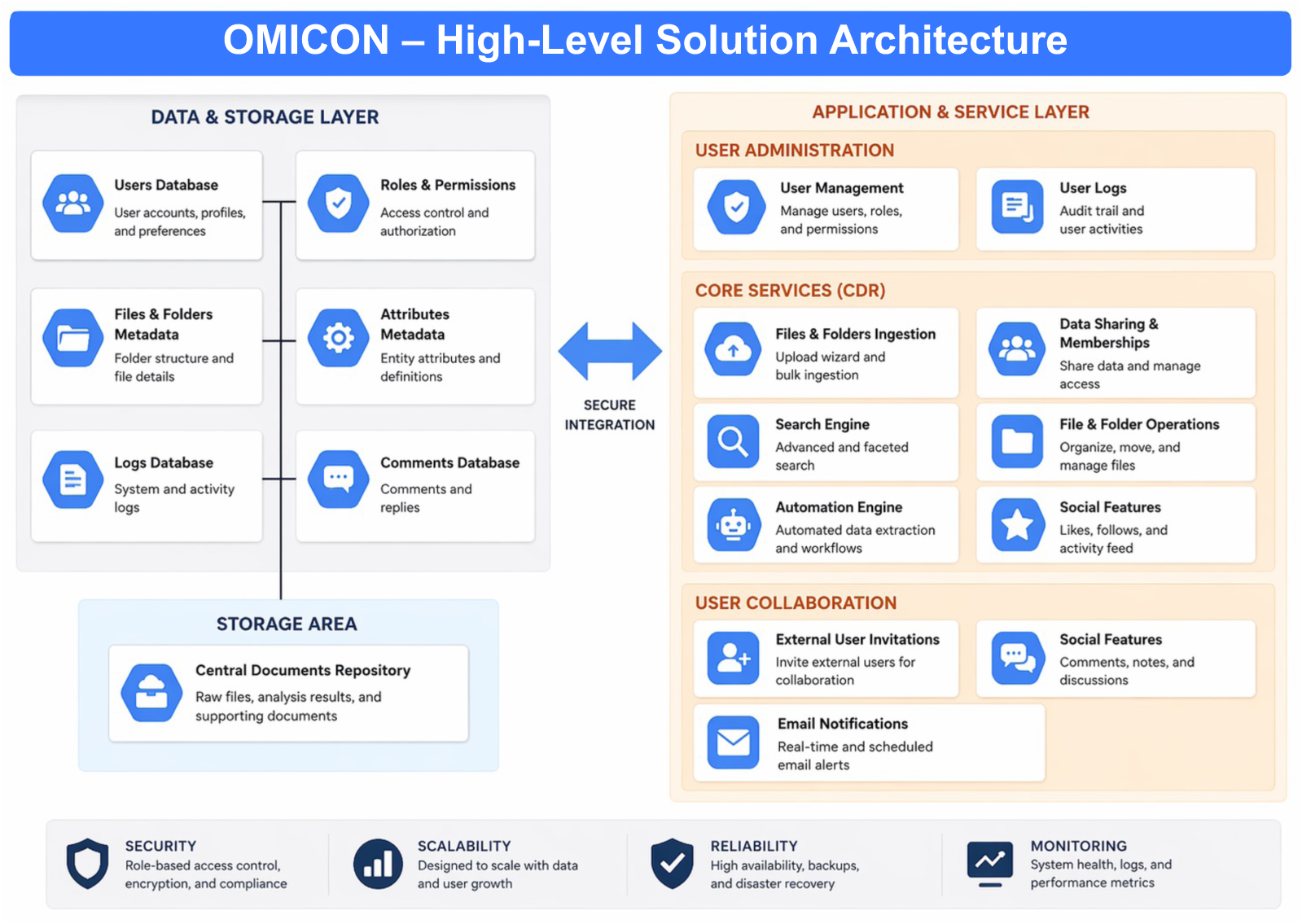
OMICON solution architecture. Overview of OMICON’s high-level solution architecture.

